# Structure of WT1 zinc fingers bound to its cognate DNA: Implications of the KTS insert

**DOI:** 10.1101/271577

**Authors:** Raymond K. Yengo, Elmar Nurmemmedov, Marjolein M.G.M. Thunnissen

## Abstract

WT1 is a transcription factor with a DNA binding N-terminal domain containing four C2H2-type zinc fingers. In order to perform its role as a transcription factor, WT1 needs to specifically recognize and properly bind to its target DNA. How this is done is still not completely clear. Two of WT1’s major isoforms are distinguished by the presence or absence of a 3 amino acid insert, Lysine-Threonine-Serine (KTS) in the linker between zinc-fingers 3 and 4. This KTS insert is conserved throughout all life forms expressing WT1. The –KTS isoform, which acts as a transcription factor, binds DNA with higher affinity than the +KTS isoform, which is thought to participate in RNA splicing and interaction with partner proteins. This study was aims at elucidating the effect of the KTS insert on DNA binding. Here we present the crystal structure of WT1 zinc fingers 2-4, with and without the KTS insert, bound to the WT1 9-base pair cognate DNA sequence, refined to 1.9 Å and 2.5 Å respectively. The structures show that the +KTS isoform of WT1 recognizes DNA with the same specificity as the –KTS isoform. The only differences in the DNA bound conformation of the two isoforms are found within the linker containing the KTS, and these mainly involve the loss of the C-capping interactions thought to stabilize the complex. These structures provide the molecular detail necessary for the interpretation of the WT1 transcriptional DNA recognition and validation of its transcriptional targets.

## Introduction

The *WT1* gene was first cloned in 1990 as a zinc finger transcription factor and RNA-binding protein that is implicated in organ development as well as tumorigenesis [1]. WT1 is known to function both as a tumor suppressor and oncogene, but the reasons behind these opposing functions are still a matter of debate. WT1 can switch roles depending on the cellular context, a phenomenon that is described by the variety of its isoforms, post-translational modifications and multiple binding partners [2, 3]. Mutations in *WT1* gene have been implicated in abnormal development of the genito-urinary system resulting in syndromes such as Wilm’s Tumor, Denys-Drash, WAGR and Frasier [4-8]. Recent studies however suggest an even wider spectrum of functions for WT1, including epigenetic regulation of genes [9-12], stem cell differentiation [13] and genome stability [14, 15].

Despite extensive studies addressing the basic biology of WT1, it is still puzzling how this protein with four C2H2 zinc fingers achieves specificity to regulate a large number of target genes involved in diverse physiological processes. To perform its transcriptional and regulatory functions, WT1 interacts with DNA, RNA and even other proteins [16-18]. Most of these promiscuous interactions occur via its zinc finger domain. Typically, each zinc finger is capable of recognizing three base pairs of DNA, although a fourth base pair binding site is preferred. The DNA binding sequence of WT1 can be summarized in the following consensus sequence 5′-GCG-(T/G)GG-G(A/C)G-(T/G)(T/A/G)(T/G)-3′ [19-26]. The non-canonical zinc finger 1 differs from the other three by its unusual amino acid composition, which partially explains the lack of stringency in the DNA binding sequence for this particular zinc finger [24]. Whereas fingers 2–4 mediate sequence-specific recognition of DNA, finger 1 does not bind DNA specifically but contributes to binding affinity [19, 23, 27].

There exist as many as 36 different WT1 isoforms, which are products of alternative splicing, alternative start codons and RNA editing. However, the four major isoforms arise due to exon 5 and exon 9. The latter encode a tri-peptide KTS, between zinc finger 3 and 4, which is more conserved among all vertebrates [6-8, 18]. Mice expressing only the +KTS or only the -KTS isoforms die shortly after birth as opposed to mice lacking the other isoforms. This indicates that the ±KTS variant is the most biologically relevant isoform [28]. Even though the expression pattern of WT1 is dependent on tissue and growth stage, a healthy ratio of +KTS:-KTS isoforms is constant at about 2:1 [29-31]. Thus, the +KTS isoform assumes more biological significance than the -KTS counterpart.

Some studies have shown that the +KTS isoform of WT1 does not bind DNA at all, while the others have shown that this isoform binds DNA very weakly [32, 33]. This reduction or abrogation of DNA binding has been attributed to the increased flexibility of the linker between zinc finger 3 and 4. The insertion of the tri-peptide KTS in this linker eliminates the helix capping mechanism that locks the bound zinc finger on the DNA or displaces zinc finger 4 from its binding site on the DNA [6, 28, 34]. A study from our lab has shown that the +KTS isoform of WT1 binds DNA with comparable affinities to the –KTS isoform, i.e in the nM range [24]. This puts in doubt the proposed hypothesis of DNA binding abrogation by KTS insertion. It is therefore still unclear how the KTS insertion affects DNA binding.

In this report we employ the structural method of X-ray crystallography to probe the molecular interaction of the WT1 zinc finger domain with its consensus DNA. Two truncations of the zinc finger domain of WT1 and a consensus DNA binding sequence were used. The resulting structures of two complexes provide molecular detail on the effect of the KTS insert on DNA binding. The structures of the last three zinc fingers of WT1 with and without the KTS insert bound to the consensus DNA sequence show that both isoforms of WT1 are capable of recognizing DNA in more or less the same manner mostly making the same interactions. This has major implications on the transcriptional role of the two major isoforms of WT1.

## Materials and methods

### Cloning, expression and purification of the WT1 ZF truncations

Four truncations of the WT1 C-terminal domain were used in this study. These included ZF24-, ZF24+ and ZF14- and ZF14+. The numbers 24 and 14 indicate the presence of zinc fingers two to four and one to four respectively. The + and – signs indicate the presence or absence of the KTS insert (**Figure 1**). The cloning of all the 4 truncations used in this study was as described previously with the exception that the expression vector used to express all 4 constructs was in this case the pET14b plasmid instead of the pETM-11, pETM30 and PCDF-1b plasmids [35].

**Figure 1.**
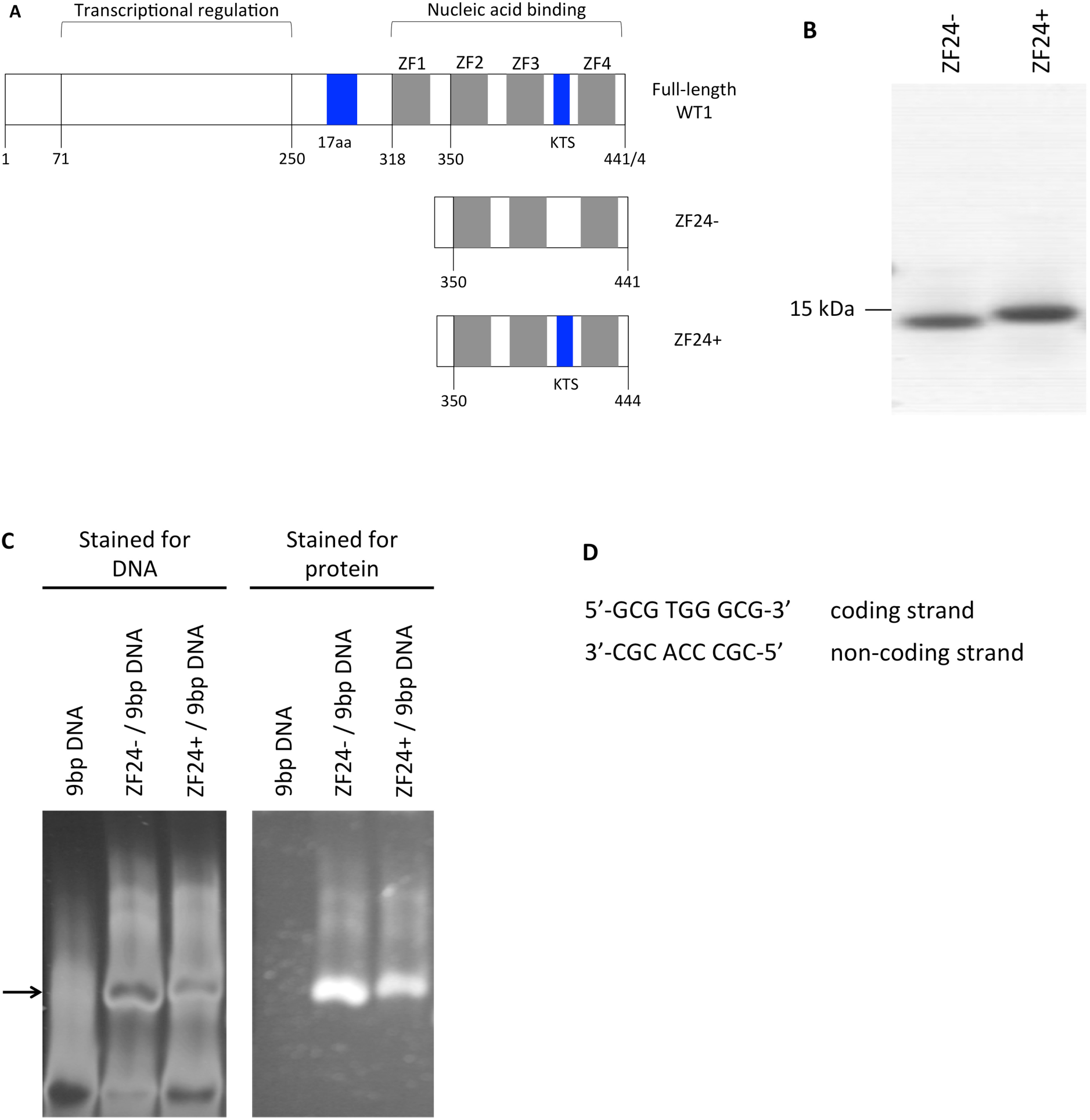
Functional and chemical properties of the components used in this study. (a) Representation of full-length WT1 and the two truncations (ZF24- and ZD24+) used in the study. (b) An SDS page gel showing the purified constructs; + and –indicate the presence or absence of the KTS insert. (c) Electrophoretic mobility shift assay showing the interaction of the ZF24- and ZF24+ with the 9bp double-stranded DNA. The gel was stained for DNA using SYBR Gold stain and for protein using SYPRO Ruby stain. The arrow indicates the zinc finger-DNA complex. (d) The DNA sequences used in the study.

The DNA plasmids pET14b-ZF24+, pET14b-ZF24-, pET14b-ZF14+ and pET14b-ZF14-were transformed into competent *E. coli* BL21(DE3)plysS cells, from which a single colony was used to inoculate 20 mL of Luria Bertani (LB) medium containing 100 µM ampicillin and 35 µM Chloramphenicol. This was grown overnight and then transferred to a 3 L flask containing 1L of LB each supplemented with appropriate antibiotics. All cultures were grown to OD_600_ ∼0.6 and induced with IPTG (0.75 mM). ZnCl_2_ was added to a final concentration of 400 µM and the cultures further incubated for 5-7 h at 37 °C to produce soluble protein. Cells were harvested by centrifugation at 5000 g for 15 min. The pellet was frozen at −20°C until purification.

The frozen pellet was thawed and re-suspended in lysis buffer (50 mM HEPES, pH 7.5, 300 mM NaCl, 2 mM MgCl_2_, 50 µM ZnCl_2_, 2 mM DTT and 1 mM PMSF), and lysed by sonication. The soluble lysate was recovered by centrifugation at 30,000 g for 30 min at 4°C. The lysate was diluted 3X and loaded onto a HiTrap SP HP column (GE Healthcare) pre-equilibrated with buffer (20 mM HEPES, pH 7.5, 10 µM ZnCl_2,_ 1mM MgCl_2_, 1 mM DTT and 1 mM PMSF). The column was washed with the same buffer containing 400 mM NaCl and the protein eluted using a linear NaCl gradient from 400 mM to 1 M. The fractions containing the protein of interest were pooled, concentrated and loaded on a Superdex 75 column (GE Healthcare) pre-equilibrated with 20 mM Tris-HCl, pH 7.0, 150 mM KCl, 10 µM ZnCl_2_, 1 mM MgCl_2_, 1 mM DTT and 1mM PMSF. The eluted protein was concentrated by ultrafiltration using Amicon Ultra centrifugal filter devices (Millipore), aliquoted, quantified and stored at −80°C until used. Analysis of the purified protein using SDS-PAGE and Dynamic light scattering showed that the proteins were homogenous with a purity of >95% (**Figure 1**). The total yield of pure homogenous protein per liter for each of the 3 constructs was 20-30 mg.

### Complex formation

The two pairs of single-stranded oligonucleotides were chemically synthesized by TAG Copenhagen A/S and shipped as lyophilized reverse phase cartridge purified DNA. Each oligonucleotide was re-suspended in H_2_O to a final concentration of 800 µM. The complementary oligonucleotides were mix in a 1:1 molar ratio, heated to 95°C for 10 min and slowly cooled to room temperature to obtain 400 µM of double stranded DNA. The DNA duplex was loaded onto a HiTrap Q FF column (GE Healthcare) pre-equilibrated in 10 mM Tris buffer (pH 7.5), and eluted with a linear NaCl gradient to separate the duplex from any extra single stranded DNA in the annealing mixture.

To obtain the complex, double stranded DNA was gradually titrated into the protein mixture to a final protein:DNA stiochiometry of 1:1.2 in binding buffer containing 20 mM Tris-HCl pH 7.5, 150 mM NaCl, 1mM MgCl_2_, 10 µM ZnCl_2_, 1 mM DTT and 1 mM PMSF. NaCl was further added to a final concentration of 500 mM to solubilise the complex. The complex was then concentrated and purified on a Superdex 75 gel filtration column in the same binding buffer containing 500 mM NaCl. The fractions containing the complex were filter concentrated, quantified and used for crystallization.

### Electrophoretic mobility shift assay

Electrophoretic mobility shift assay (EMSA) was performed to ascertain the DNA binding activity of the different WT1 zinc finger truncations. 1ug of a 12bp DNA probe shown to be recognized by WT1 was incubated with equivalent molar amounts of all 9 WT1 zinc finger truncations in our lab in buffer containing 20 mM Tris-HCl pH 7.5, 150 mM KCl, 2 mM MgCl_2_ 2 mM DTT, 2 µM ZnCl_2_ and 50% Glycerol. The reaction was incubated at room temperature for 15 – 20 min. Separation of the reaction mixture was carried out on an 8% nondenaturing polyacrylamide TB gel. The gel was stained with the nucleic acid stain SYBR Green and subsequently with the protein stain SYPBR Ruby from the Electrophoretic Mobility Shift Asssay Kit (E33075) from Invitrogen and visualized on a UV transilluminator.

### Crystallization, structure determination and analyses

Hexagonal rod-shaped crystals were grown by the hanging drop vapor diffusion method at 20°C. For the ZF24-9bp structure, the drop contained 1 µL of complex and 1 µL of reservoir solution consisting of 50 mM HEPES pH 7.5, 15 mM MgCl_2_, 1.0 mM Spermidine and 10 % Dioxan. For the ZF24+9bp structure the drop contained 1 µL of complex and 1 µL of reservoir solution consisting of 50 mM Cacodylate pH7.0, 20 mM MgCl_2_, 2.5 mM Spermine and 10 % Isopropanol. Crystals appeared after 2-4 days and grew to their full size in one to two weeks. The crystals were transferred to a cryo solution containing the reservoir solution and 25% Glycerol and flash cooled in liquid nitrogen. Data was collected at the Cassiopeia I911-3 beamline at the MAX-lab synchrotron radiation facility in Lund, Sweden. The diffraction data was integrated, scaled and converted to structure factors using the XDS data processing package [36]. The ZF24-9bp structure was solved using a combination SAD (Single Anomalous Scattering) and MR (Molecular replacement). The position of the 3 zinc atoms was located using the SHELX program package through the HKL2MAP interphase [37] and the initial phases used in the molecular replacement solution program PHASER from CCP4 [38] using the 1AAY structure as search model. An initial model was built into the electron density map manually in Coot [39]. The ZF24+9bp structure was solved by molecular replacement with Phaser using the ZF24-9bp structure as search model. Rigid body refinement was followed by multiple steps of TLS and restrained refinement with Refmac from the CCP4 suite [40]. The TLS groups were chosen such that each zinc finger, DNA strand and the zinc atoms formed a unique group. Evaluation of model fit to density, model building and correction as well as the addition of waters was performed using Coot [39]. The final structure refinements were carried out using Phenix refine [41]. Molecular graphic figures were prepared with PyMOL [42].

## Results

### Protein purification and electrophoretic mobility shift assay (EMSA)

The WT1 zinc finger truncations used in this study were purified using a combination of ion exchange chromatography and gel filtration. The purified proteins had a purity of 90% or more as can be seen in **Figure 1**. The homogeneity of the samples were verified by dynamic light scattering and native gel electrophoresis and the samples were all homogenous even after storage at −80°C as concentrated stock solutions of up to 60 mg/ml for 1 month. An EMSA was run to ascertain that the samples were correctly folded and active. The EMSA native gels were first stained for DNA and then for protein. The first stain (SYBR Green) shows the position of the complex bands with respect to the DNA probe and the second stain (SYPRO Ruby) shows the position of the complex since the protein is not seen because it migrates in the opposite direction due to its high pI. The stained gels (**Figure 1**) show that both the –KTS and +KTS isoforms of WT1 bind DNA with high enough affinities to be separated on native gel.

### Structures of ZF24-bound to WT1 cognate 9 bp DNA (ZF24-9bp)

A truncation of the WT1 zinc finger domain including fingers 2-4 (residues 350-441) with an N-terminal methionine introduced by the cloning method and excluding the KTS insert at the linker region between zinc finger 3 and 4 (between residue 407-408) purified to homogeneity was used for crystallization. The complex between the purified protein (shown in **Figure 1**) and the WT1’s 9 bp cognate DNA sequence was crystallized after complexes with longer DNA up to 13 bp failed to crystallize. Data was collected on the hexagonal crystals, which grew to their full size in about 2 weeks and indexed using the XDS program package in space group P6(5) (**Table 1**). The phases were obtained by a combination of single anomalous dispersion [43] and molecular replacement [44] with 1 molecule per asymmetric unit and a solvent content of 46.6%, using the structure of Zif268 in complex with DNA (PDB code 1AAY) as a search model. Amino acid residues 352 to 437 were built and refined but electron density could not be observed for the first 2 and last 4 residues most likely due to disorder. Refinement of the structure to 2.5 Å enabled all the DNA nucleotides to be correctly modeled but only 4 water molecules could be modeled. The final, refined model (**Figure 2**) has an R_work_ of 18.8%, R_free_ of 26.2, a B-factor of 56.93 and all backbone φ and ψ angles are within allowed regions. A full summary of the data collection and refinement statistics is shown in **Table 1**.

**Table 1:** Data collection and refinement statistics Values in parentheses are for the highest resolution shell.

|  | **ZF24-9bp** | **ZF24+9bp** |
| --- | --- | --- |
| Data collection |  |  |
| Beamline (Å) | Maxlab I911-3 | Maxlab I911-3 |
| Wavelength | 1.283 | 0.908 |
| Resolution | 25-2.5 (2.6-2.5) | 30-1.9 (2.0-1.9) |
| Space group | P 6(5) | P 6(5) |
| Unit-cell parameters | a=b=39.45 Å, c=173.61 Å<br>$\alpha=\beta=90.00^\circ$, $\gamma=120.0^\circ$ | a=b=37.13 Å, c=173.16 Å<br>$\alpha=\beta=90.00^\circ$, $\gamma=120.0^\circ$ |
| Measured reflections | 40161 (5173) | 77384 (11784) |
| Unique reflections | 9840 (1353) | 20606 (3200) |
| Redundancy | 4.08 (3.82) | 3.76 (3.68) |
| Completeness | 93.6 (80.1) | 97.4 (93.4) |
| $\langle I/\sigma(I) \rangle$ | 16.81 (4.16) | 19.9 (6.5) |
| R<sub>merge</sub> (%) | 7.8 (43.4) | 5.3 (21.0) |
| R<sub>meas</sub> (%) | 6.0 (34.9) | 5.2 (23.5) |
| Anomalous correlation (%) | 47 (6) | - |
| Refinement |  |  |
| Reflections (total) | 5243 | 10022 |
| Reflections (test-set) | 263 | 528 |
| Resolution range | 20-2.5 | 25-1.9 |
| R<sub>work</sub> (%) | 18.8 (23.9) | 16.9 (24.5) |
| R<sub>free</sub> (%) | 26.2 (31.9) | 23.1 (27.1) |
| Residues/Protein atoms | 86/735 | 90/764 |
| Nucleotides/DNA atoms | 18/369 | 18/369 |
| Metal atoms (Zn) | 3 | 3 |
| Solvent atoms | 4 | 149 |
| Wilson B (overall) | 56.93 | 26 |
| B value (average) | 64.9 | 35.4 |
| RMSD from ideal geometry |  |  |
| Bond lengths (Å) | 0.013 | 0.007 |
| Bond angles (°) | 1.265 | 1.028 |
| Ramachandran (%) |  |  |
| Favoured | 90.5 | 94.3 |
| Allowed | 9.5 | 5.7 |

**Figure 2.**
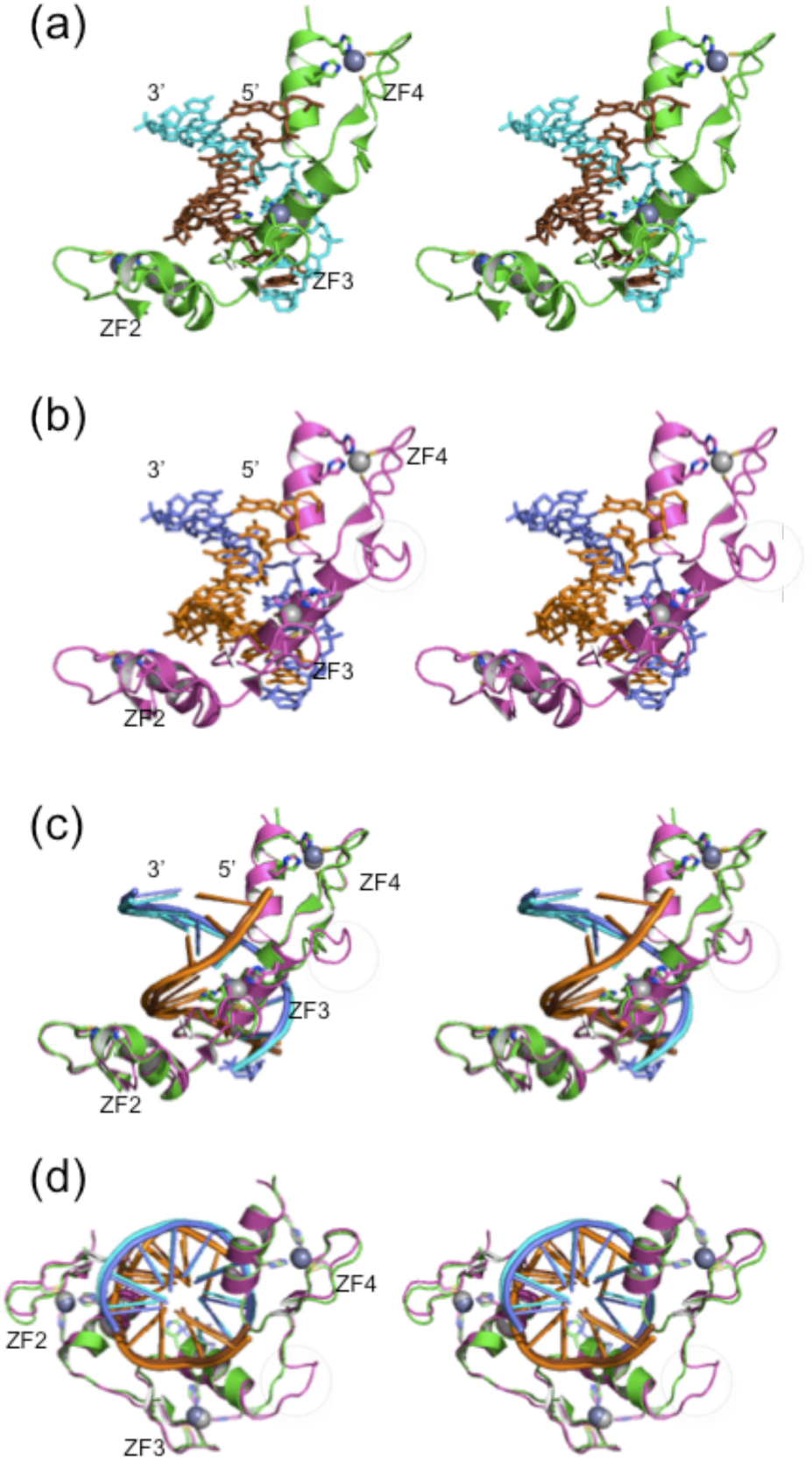
Stereo view of the individual structures and superposition of the two structures. (a) Structure of ZF24-(in green) bound to a 9 bp DNA (ZF24-9bp). The DNA coding strand is in brown and the non-coding strand in cyan. (b) Structure of ZF24+ (in magenta) bound to a 9bp DNA (ZF24+9bp). The DNA coding strand is in orange and the non-coding strand in slate. (c) The superposed ZF24-9bp and ZF24+9bp structures viewed sideways. (d) The superposed ZF24-9bp and ZF24+9bp structures viewed from above.

In the ZF24-9bp structure, the zinc fingers recognize the DNA in the conventional manner in which C2H2 zinc fingers recognize DNA wrapping around the DNA double helix with the α-helix from each finger, packing against the major groove. All the 3 zinc fingers have well defined electron density showing well defined C2H2 zinc fingers consisting of a β hairpin loop and an α-helix held together by a Zn^2+^ ion coordinated by a pair of cysteines and a pair of histidine residues. The fingers are docked on the DNA in such a way that the α helices lie at an angle of 57.8° between finger 2 and 3 helices and 68.1° between fingers 3 and 4 helices. The bound DNA is a modified version of B-form DNA typical of zinc finger DNA interactions. The major groove of the DNA in this structure is wider and a bit deeper than that of an ideal B-form DNA. A structure superposition of an idealized B-form DNA and the DNA in this structure shows a backbone atom RMSD of 3.4 Å. This difference is mainly as a result of the partial unwinding of the DNA double helix to accommodate the interacting α-helices from the protein.

In this structure the DNA is involved in crystal packing, packing tail-to-head with G1 packing against G9 and C59 packing against C51. The distance between the last base-pair from one molecule and the first base-pair in the next molecule is an ideal stacking distance for B-form DNA. The packing also respect the strands with the primary strand packing tail to head with another primary strand such that the crystal seems to be made up of long continuous DNA with a long continuous string of zinc fingers wrapped around it. In the diffraction images collected from the crystals this arrangement was already visible since a characteristic DNA fiber diffraction pattern was observed. In addition, another interesting crystal packing interaction depicted in **Figure 4**, also occurs at the tip of the structure with finger 4 lying at the crystal packing interface in a major groove contributed by two halve DNA sites from different complexes. The finger does not make any specific hydrogen bond interactions with the DNA half site from the next complex but it makes a stabilizing hydrogen bond with the backbone phosphate of G9 via the side-chain of HIS431.

**Figure 3.**
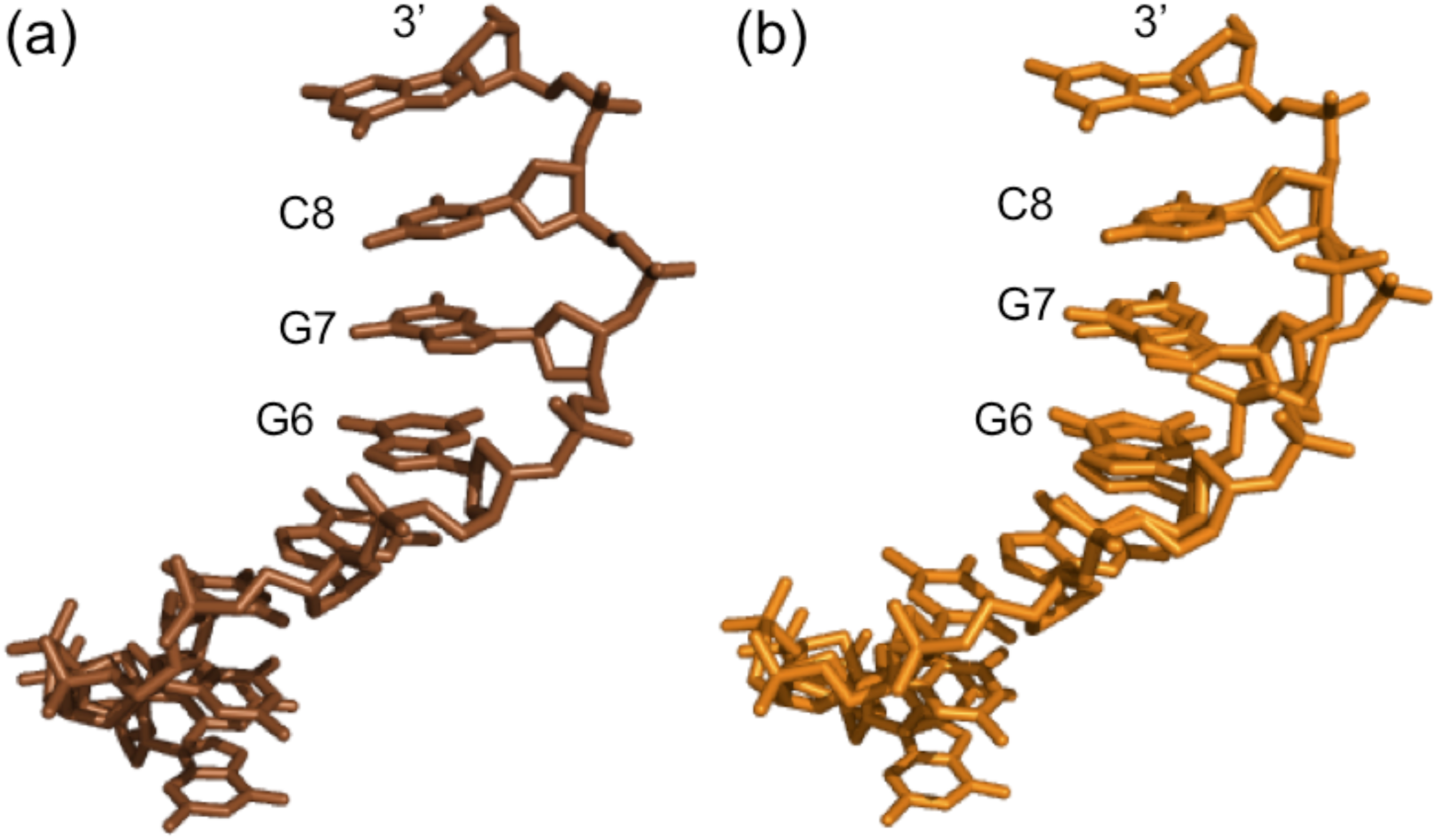
Comparison of the DNA structures in ZF24-9bp and ZF24+9bp complexes. (a) The B form undistorted coding strand in the ZF24-9bp structure. (b) The distorted backbone conformation of the coding strand in the ZF24+9bp structure modeled in a double conformation.

**Figure 4.**
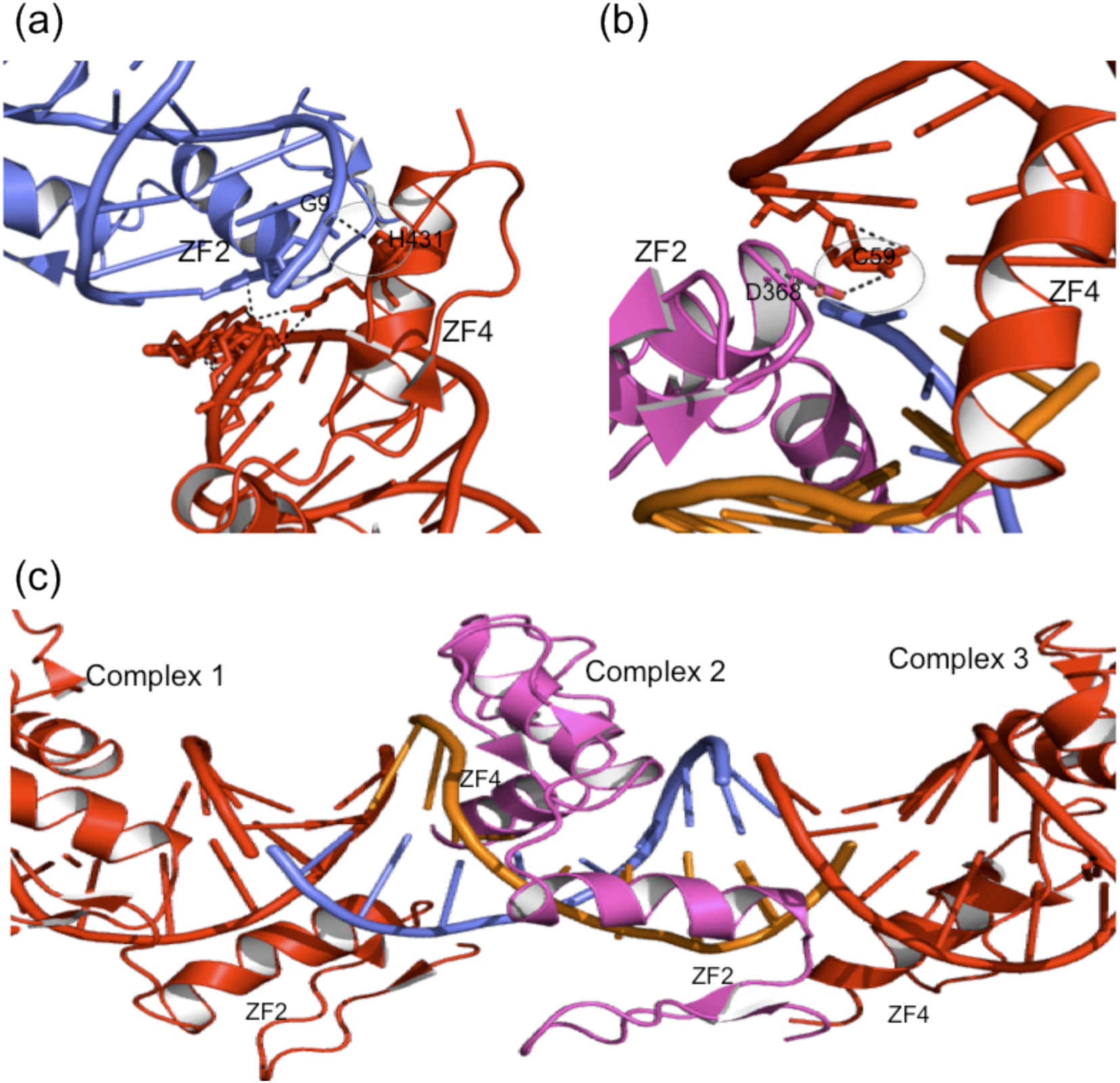
Crystal-packing interactions in ZF24-9bp and ZF24+9bp structures. (a) In the ZF24-9bp structure, the α-helix in zinc finger 4 lies in the major groove contributed by DNA from adjacent molecules making a backbone hydrogen bond interaction with the adjacent DNA. (b) In the ZF24+9bp structure, Asp368 at helix position 1, makes a base specific H-bond with G59 from the adjacent molecule. (c) The DNA packs head to tail with the terminal base pairs stacking to create a long disconnected double helical DNA with a tandem array of disconnected 3 zinc finger repeats wrapped around it.

### Structure of ZF24+ bound to WT1 cognate 9 bp DNA (ZF24+9bp)

The same construct as for the ZF24-was used, but for the fact that this one includes the KTS insert (residue 408-410), which results in the re-numbering of the downstream sequence (residues 350-444). The structure of the complex between the purified protein (also shown in **Figure 1**) and the WT1 9 bp cognate DNA was determined. Data was collected on the crystal, which was also hexagonal, though grown under different conditions and indexed and processed using the XDS program package. The space group for these crystals was P6_5_ (**Table 1**). Phases were obtained by using molecular replacement for one molecule per asymmetric unit and a solvent content of 46.6 %, using the ZF24-9bp structure as a search model. Amino acid residues 351 to 439 were built and refined but no electron density could be observed for the first and last 5 residues. Refinement of the structure to 1.9 Å enabled the modeling of all the DNA nucleotides and 149 water molecules. The final, refined model represented in **Figure 2**, has an R_work_ of 16.9%, R_free_ of 23.1, a B-factor of 26 and all backbone φ and ψ angels are within allowed regions. A full summary of the data collection and refinement statistics is shown in **Table 1**.

The ZF24+9bp structure is almost identical to the ZF24-9bp structure with the only immediately apparent difference being the protruding structure formed by the KTS insert (**Figure 2**) and the double conformation of the DNA (shown in **Figure 3**). The RMSD between these two structures of all backbone atoms is 0.94 Å and of all atoms is 1.43 Å. The inter-finger interactions represented in **Figure 5** are identical in both structures. Even the crystal packing (**Figure 4**) interactions in the ZF24+9bp structure are very similar to those in the ZF24-9bp structure but for the packing interaction mediated by the KTS insert and the extra hydrogen bond interactions, which is a base specific interaction between ASP368 in zinc finger 2 and C59 from the preceding complex molecule.

**Figure 5.**
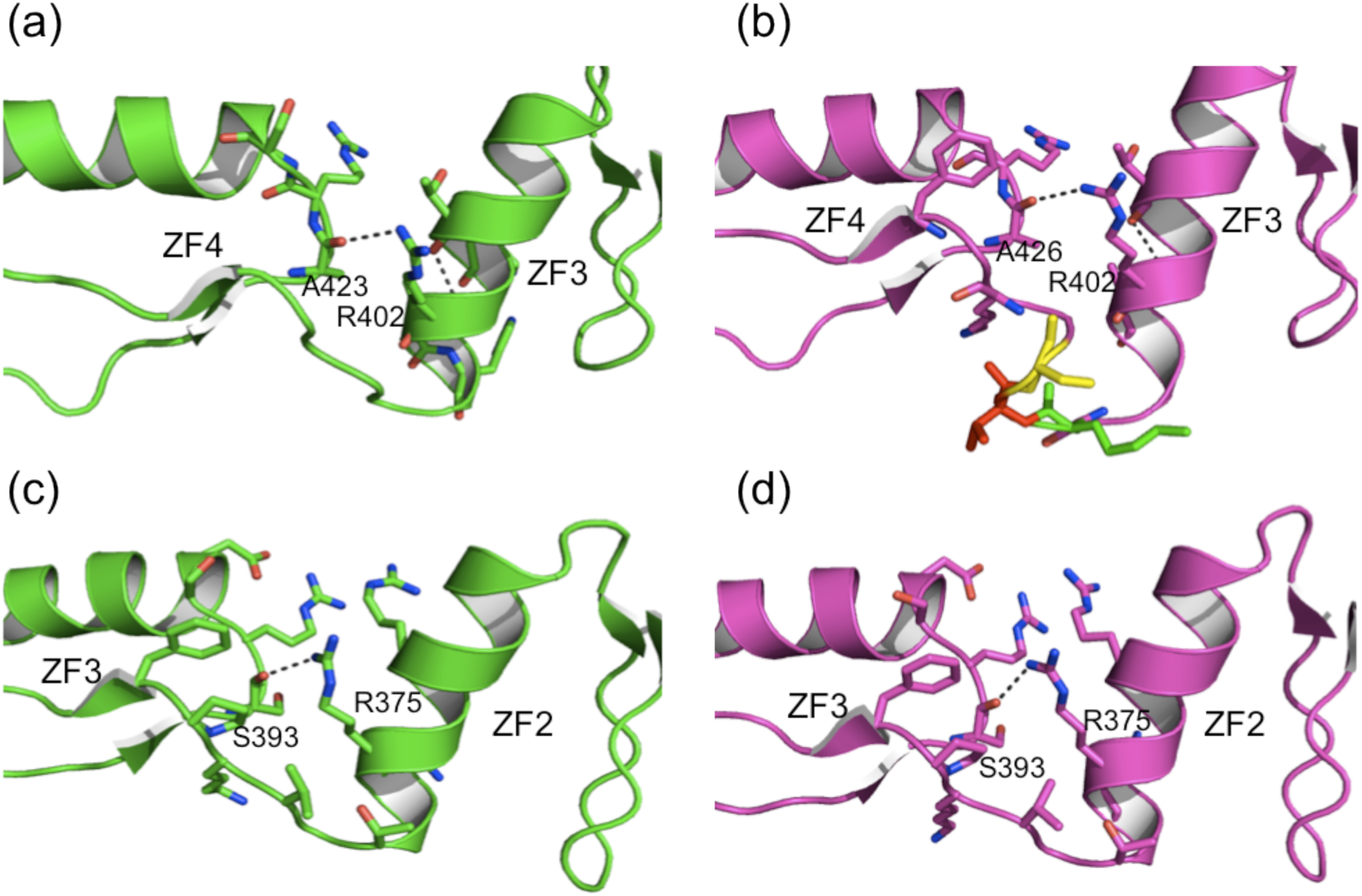
Inter-finger interactions between adjacent fingers. The inter finger interactions between ZF4 and ZF3 are identical in both the ZF24-9bp structure (a), and the ZF24+9bp structure (b), despite insertion of the KTS insert at this interface. The inter finger interactions between ZF3 and ZF4 are identical in both the ZF24-9bp structure (c), and the ZF24+9bp structure (d) as expected.

### Individual zinc finger interactions with DNA

C2H2 zinc fingers conventionally recognize DNA using specific amino acid positions within the α helix. The hydrogen bond interactions mediated by these amino acid positions that define the specificity of WT1 for its target DNA are illustrated in **Figure 6** while **Figure 7** present a summary of all the interactions made by WT1 zinc finger 2 to 4 in DNA binding. The specific interactions mediated by zinc finger 2 are identical in the ZF24-9bp and the ZF24+9bp structures. These base-specific interactions from zinc finger 2, as observed in these structures include a hydrogen bond made between Arg372 and G6 and the hydrogen bond between Gln 369 and C8. The only difference in this interaction between the two structures is that Arg372 make two hydrogen bonds with G6 in the ZF24+9bp structure compared to only one in the ZF24-9bp structure. Three amino acids from zinc finger 3 are involved in base specific interactions in both structures. Arg394 makes two hydrogen bonds with G7, Asp396 makes one hydrogen bond with C53 and His397 makes one hydrogen bond with G5. The only noticeable difference here is the additional water-mediated interaction between ASP396 and C54 in the ZF24+9bp structure, which is understandably absent in the ZF24-9bp structure since due to resolution limitations of the data, considerably few waters molecules were modeled. Two amino acids from zinc finger 4 make base specific hydrogen bonds with the DNA. ARG 427(Arg424) makes two identical hydrogen bonds with G3 in both structures (numbering in parentheses reflect the –KTS isoform). Arg433(ARG430) makes a hydrogen bond with G1 and another with G58 in the ZF24-9bp structure. The same Arg433(ARG430) however makes two hydrogen bonds in the ZF24+9bp structure, one with G1 and the other, a water mediated interaction, with G58. Overall, the base specific interactions made in the two structures is mostly identical but for a few minor differences which will mostly influence the binding affinity rather than binding specificity.

**Figure 6.**
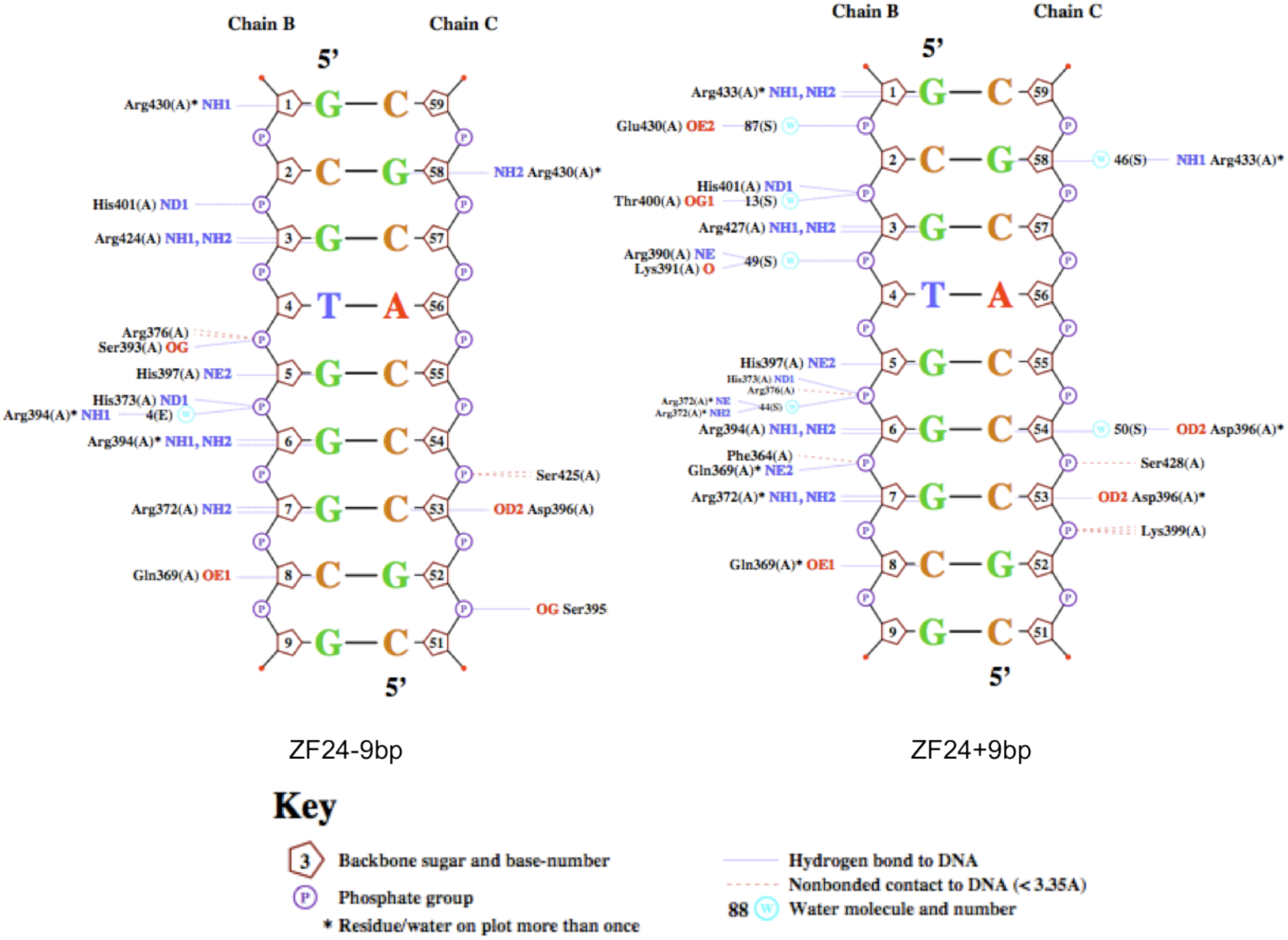
An overview of the interactions in ZF24-9bp and ZF24+9bp structures calculated by Nucplot. Amino acid residue numbering is the same for zinc finger 2 and 3. Amino acid 408-437 in the ZF24-9bp structure corresponds to amino acid 411-440 in the ZF24+9bp structure due to insertion of the KTS insert between amino acid 407 and 408.

**Figure 7.**
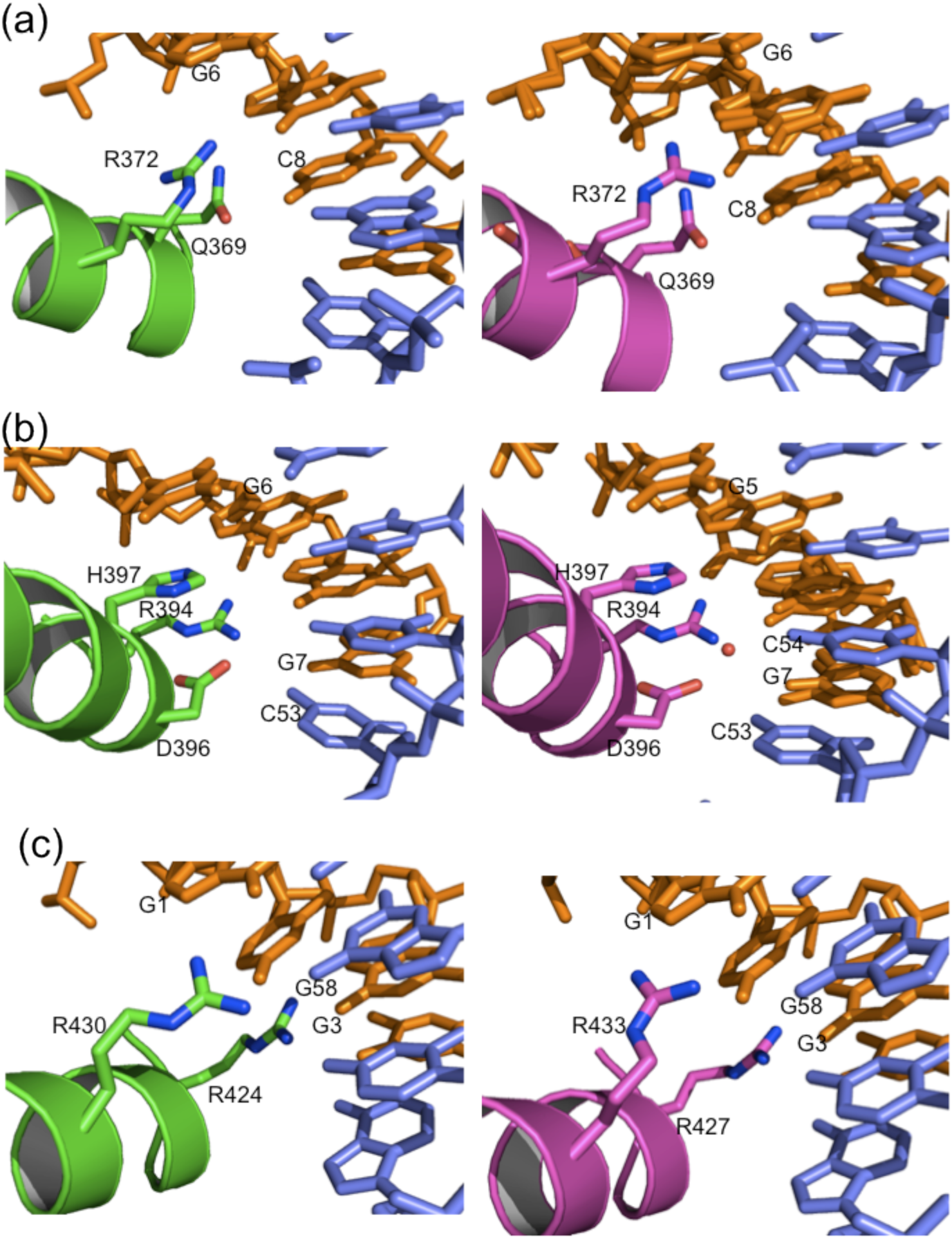
Comparison of the base specific interactions mediated by the individual zinc fingers in the ZF24-9bp structure (left panel) and the ZF24+9bp structure (right panel). The color code is same as in figure 2. (a) Finger 2, (b) Finger 3 and (c) finger 3.

### Linkers

All of the linkers in the structure of ZF24-bound to the 9 bp cognate DNA adopt well-defined conformations (**Figure 8**). The linkers pack at the finger interface with one surface against the interface created by the α-helix of the preceding zinc finger and the N-terminal tip of the following zinc finger. The packing interactions mediated by the linkers at the finger interface are predominantly backbone mediated hydrophilic interactions. The side-chains of the linker are arranged to the side of the interface packing against each other in a hydrophobic environment to provide a stable structure. The interface between zinc finger 2 and 3, and that between zinc finger 3 and 4 are almost identical. In both cases the linker is anchored in place by two hydrogen bonds, one at each end. The anchoring hydrogen bond at the C-terminal end is made between the backbone carbonyl oxygen of Phe 392 and Phe422 to the backbone amide of Phe383 and Phe411 in the zinc finger 2 and 3 interface and zinc finger 3 and 4 interface respectively. The anchoring hydrogen bonds at the N-terminal end of both the zinc finger 2 to 3 and 3 to 4 linkers are the α-helix capping hydrogen bonds. These hydrogen bonds are made between the amide of the conserved Gly379 and Gly407 following zinc fingers 2 and 3 respectively and the backbone carbonyl of the third to last residue on the preceding α-helix [45]. There is no equivalent linker sequence or glycine following zinc finger 4, but since this helix capping hydrogen bond is important for the stability of the DNA binding α-helix, the arginine at the equivalent position mediates this hydrogen bond with its side-chain NH2 group. The linker is further held in place by the C-terminal α-helix Thr378 and Thr406 of helix 2 and 3, which hydrogen-bonds to the backbone carbonyl of Glu374 and Thr402 respectively via its backbone amide as well as to the backbone amide of the second linker residue via its side-chain carbonyl. The two interfaces between zinc fingers 2 and 3 and zinc fingers 3 and 4 are stabilized by Arg375 and Arg403, which makes hydrogen bond contacts with the backbone carbonyl of ARG394 and Arg424 respectively both at helix position −2 via its side chain NH2. Arg403 additionally makes a hydrogen bond via its side-chain amine with the side-chain carbonyl of Glu408 completing an intricate and well-conserved hydrogen bond network designed to maintain a particular interface between C2H2 zinc fingers in DNA binding domains.

**Figure 8.**
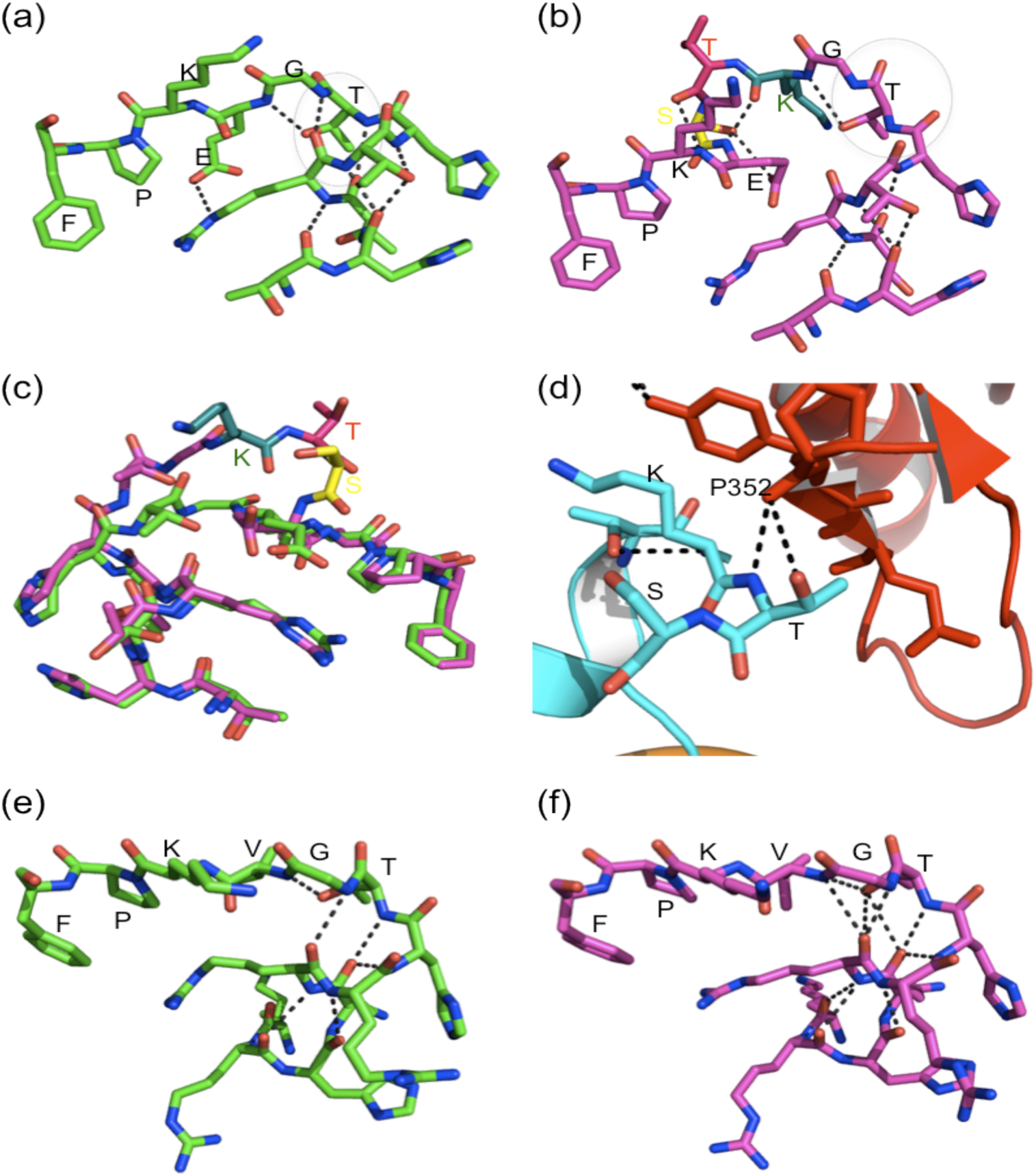
Comparison of the conformation and interactions made by the linkers between the zinc fingers in ZF24-9bp and ZF24+9bp structures. (a) and (b) show the conformation and interactions in the linker between ZF3 and ZF4 in the ZF24-9bp structure and the ZF24+9bp structures respectively. The helix capping interactions mediated by glycine and threonine in the ZF24-9bp structure are lost in the ZF24+9bp structure. Superposition of this linker from the two structures in (c) show their differing conformation. (d) shows the crystal packing interaction made by lysine which is part of the KTS insert in the ZF24+9bp structure. (e) and (f) show the comparison of the linker between ZF2 and ZF3 in the ZF24-9bp structure and the ZF24+9bp structure respectively. The linker in the ZF24+9bp structure makes all the interactions like in the ZF24-9bp structure but also makes a number of additional interactions.

In the structure of ZF24+ bound to the cognate 9 bp DNA the linkers also adopt well-defined structural conformations. The structure and the interactions responsible for the interface between zinc finger 2 and 3 is identical to that in the structure ZF24-9bp structure but for the fact that the angle between the linker and the preceding helix is a little smaller. This reduced angle results in a more extensive hydrogen bond network that cap the helix even more firmly. There are however, major differences between the linker conformation and the interface interactions between zinc finger 3 and 4. These differences are to be expected given the sequence variation between the linkers in the two structures. The threonin and serine from the KTS insert together with the normal linker glutamate folds into a single turn of a 3_10_ helix. The backbone carbonyl of LYS408 from the KTS insert makes a hydrogen bond with the backbone amide of Glu411 while the backbone carbonyl of Thr409 makes a hydrogen bond with the backbone amide of LYS412. These two hydrogen bonds define the 3_10_ helix, which is locked in its position by a hydrogen bond formed between the side chain carbonyl of Glu411 and the side chain OH of Ser410. Lys424 loops over from the second β strand to anchor the 3_10_ helix from the C-terminal end while the conserved Thr406 anchors it from the N-terminal side. All other interface hydrogen bonds are identical to those made in the ZF24-9bp structure with the exception that the helix capping hydrogen bonds mediated by the conserved linker glycine and the terminal helix amino acid, threonine are notably missing. This loss of the α-helix capping interactions has been earlier proposed [32]. The KTS insert in this structure makes a crystal packing interaction with Thr409 making two hydrogen bonds with the backbone carbonyl of Pro352 via its side chain carbonyl and backbone amide.

## Discussion

The general principle by which WT1 recognizes DNA in the two structures presented here is analogous to that presented earlier [27, 32]. Very minute differences are observed between the ZF24-9bp structure and the previously reported structures of WT1 and Zif268 [46]. Whereas in the previous structures only a proposal for the hydrogen bonding interactions could be given due to limited resolution, the structures reported here allow for a more detailed description. The ZF24+9bp structure on the other hand presents a first look at the molecular interactions that govern the differential recognition of DNA by the +KTS and the –KTS isoforms of WT1. It clearly shows the reason for the reasonably stable complex between the WT1 zinc fingers and its cognate DNA, as demonstrated in the EMSA experiment and also in previous BIACORE experiments [24]. Interestingly, the base-specific interactions of both isoforms with DNA are identical and even the non-specific interactions with the DNA backbone are also mostly identical. The major differences are observed away from the DNA binding surface, around the linker region where the KTS is inserted. These mostly involve inter- and intra-zinc finger interactions, which are apparently very important in the WT1-DNA complex. The other major difference is observed at the level of the DNA conformation since the density for the DNA in the ZF24+9bp structure shows the presence of two slightly different conformations. This distortion in DNA conformation may be due to the stress exerted on the DNA by the bound protein. The DNA recognition interactions at the packing interface where the terminal zinc fingers recognize DNA nucleotides in the adjacent molecule in the crystal suggest that the WT1 DNA binding sequence may be an 11 base-pair long sequence represented by 5’-GGCGTGGGCGG-3’. The crystal packing interactions involving the DNA head-to-tail packing presents a possible model for longer DNA sequence recognition by zinc fingers in tandem array.

The fact that zinc finger 4 in the ZF24+9bp structure binds DNA in an identical manner to the way it does in the ZF24-9bp structure has been suggested earlier by surface plasmon resonance (SPR) studies, which only showed a 2-fold decrease in DNA binding affinity for ZF14+KTS compared to ZF14+KTS [24]. The same study however showed a 17-fold decrease in DNA binding affinity for ZF24+KTS compared to ZF24-KTS. This noticeable difference in the latter case ascertains that zinc finger 1 plays a significant role in DNA binding elevating the affinity to compensate for the effect of the KTS insert. A recent NMR study suggests that the KTS insert prevents zinc finger 4 from optimal binding to DNA possibly by displacing it from its binding site [33]. This would suggest the formation of a rigid structure in the original linker to enable such a displacement. The NMR study however did not permit the elucidation of such structural detail. The results from the present study show in molecular detail that zinc finger 4 in the +KTS isoform binds DNA in an almost identical way as in the –KTS isoform. This contradicts the hypothesis that zinc finger 4 is displaced from its binding site by the KTS insert. Analysis of the interactions made by zinc finger 4, both base specific and backbone interactions, do not suggest any difference in binding mode or potential displacement of zinc finger 4 from it’s binding site. It is therefore plausible that the difference in binding affinity between the –KTS and +KTS isoform of WT1 is not necessarily related to the displacement of zinc finger 4 from its binding site, but is rather linked to the change in the binding energetics or some other kinetic parameters. Indeed, SPR studies show that in the case of ZF24+, the association rate remains unchanged in comparison to ZF24-, while the dissociation rate is decreased by a factor of 5. This could suggest that the KTS insertion makes the linker more flexible and zinc finger 4 therefore takes longer to find its target DNA binding site, but once bound, it make as strong contacts as the –KTS isoform, as indicated by the identical binding contacts in the ZF24-9bp and the ZF24+9bp structures.

Examination of all the linkers in the ZF24-9bp and ZF24+9bp structures show that they all adopt identical, well-defined conformations except for the linker between zinc fingers 3 and 4 in the ZF24+9bp structure that contains the KTS insert. All the other linkers adopt a stretch-out conformation typical of linkers in DNA-binding C2H2 zinc fingers with the consensus sequence G(E/V)KP(F/Y), such as the linkers in Zif268. These typical linkers all make the α-helix capping interaction mediated via the backbone amide of the first glycine in the linker and the backbone carbonyl of the last residue in the helix (threonine) to the third and forth to last residue in the preceding helix respectively [45]. This is consistent with results from other structural studies of WT1 in particular [27], and C2H2 zinc finger in general bound to DNA [46]. The linker containing the KTS insert adopts an alternative conformation in which the KTS participates in a short 3_10_ helix which folds away from the finger interface. The density observed for the linker in the crystal structure was of very high quality and the conformation of the loop could be built unambiguously. The significance of the rare 3_10_ helix adopted in this structure is unknown but its formation and special orientation increases the angle between the helix and the loop disrupting the helix capping interactions. It has been shown earlier that the KTS insertion disrupts the helix capping interactions but it was unclear why that should be the case since the Thr406 and Gly407 are still present in their original position as in the –KTS isoform [32, 34]. The special orientation of the KTS and the fact that these helix-capping residues are stretched out, away from their ideal hydrogen bonding positions, coupled with the fact that Glu408 loses its interaction with Arg403 in the ZF24+9bp structure, explains this earlier observation. The previous NMR studies also predicted that the loss of the helix capping interactions between zinc finger 3 and the linker could lead to an unstable protein unable to bind DNA. In this structure, despite the loss of the helix capping interactions, there is no major de-stabilization observed and the complex formation with DNA is not impaired.

It is still unclear why there is a reduction in DNA binding affinity of the +KTS isoform. It has been proposed that the longer linker results in an entropic gain for the free protein [32, 33]. It is thus possible that the enthalpic gain derived from the formation of the 3_10_ helix may not be enough to compensate efficiently for the entropy loss due to the binding of the longer and more flexible linker, given that some of the interactions such as C-capping normally contributing to the enthalpy gain upon binding are lost. The binding of the +KTS isoform of WT1 to unmodified DNA may therefore not be as energetically favorable as the binding of the –KTS isoform resulting in their differential DNA binding properties. The data presented here and elsewhere indicates that the difference in DNA binding affinity between the –KTS isoform and the +KTS isoform of WT1 may not be as large as originally thought [20, 32, 47]. The WT1+KTS isoform can still bind DNA with sufficient affinity, but may have preference for epigenetically modified DNA. Indeed, a recent study clearly demonstrates that ZF24+KTS has significantly higher affinity for DNA containing 5-carboxylcytosine and 5-methylcytosine than unmodified DNA [48]. The study also posits that Frasier syndrome stems from perturbed binding at genomic sites that favor binding of the –KTS isoform.

The significance of the +KTS and –KTS isoforms of WT1 in development and disease is undoubted. While there are reports of the +KTS isoform functioning as a transcription factor [49] in certain cell models, comparative investigation of the two isoforms requires a more holistic cell-based approach. A recent deep sequencing study has revealed distinct global binding patterns for the +KTS and –KTS isoforms [50]. It is shown that even though both isoforms have common gene targets, binding of -KTS is specific to transcription start sites and to enhancers, whereas binding of +KTS is more abundant and is specific to regions within gene bodies. This opens a possibility that +KTS binds DNA indirectly via recruitment of other partner proteins, and that these interactions are low-affinity transient interactions. In fact, in our crystal structure of ZF24+9bp, the threonine residue inside the KTS insert is found to make a crystal-packing contact with a neighboring zinc finger (**Figure 4**), suggesting its role in protein-protein interactions. All this is also consistent with possible involvement of KTS in epigenetic regulation as well as its role as a spacer, which would allow transient interactions with less preferred binding sites of WT1.

The present structures of the WT1 zinc fingers 2-4 bound to DNA show that the +KTS isoform can bind the same DNA sequences, with identical specificity to the –KTS isoform. We show that the differences in the molecular interactions involved in this binding exist mainly at the site of the KTS insertion with the loss of some of the stabilizing interactions in the DNA bound protein. These structures shed more light on the molecular details responsible for the differential recognition of DNA by the –KTS and +KTA isoforms of WT1. Such details will contribute to a better understanding of WT1’s functional role as a transcription factor.

